# Optimal diameter reduction ratio of acinar airways in human lungs

**DOI:** 10.1101/410522

**Authors:** Keunhwan Park, Taeho Son, Young-Jae Cho, Noo Li Jeon, Wonjung Kim, Ho- Young Kim

## Abstract

In the airway network of a human lung, the airway diameter gradually decreases through multiple branching. The diameter reduction ratio of the conducting airways that transport gases without gas exchange is 0.79, but this reduction ratio changes to 0.94 in acinar airways beyond transitional bronchioles. While the reduction in the conducting airways was previously rationalized on the basis of Murray’s law, our understanding of the design principle behind the acinar airways has been far from clear. Here we elucidate that the change in gas transfer mode is responsible for the transition in the diameter reduction ratio. The oxygen transfer rate per unit surface area is maximized at the observed geometry of acinar airways, which suggests the minimum cost for the construction and maintenance of the acinar airways. The results revitalize and extend the framework of Murray’s law over an entire human lung.

## MAIN TEXT

### Introduction

Fluid transport systems in the form of branching networks have evolved in multicellular organisms to deliver bulk metabolic matter to matter exchange sites (*1-7*). In a branching network, a mother branch is divided into numerous terminal daughters. The aggregate cross-sectional area of the vessels of a single generation generally increases with branch generations, and the flow velocity thus decreases. A low flow speed is advantageous for allowing more time for mass transfer at the terminal branches (*1*). For instance, the diameter of vascular vessels in the human body decreases from ∽ 1 cm at the aortae to ∽ 10 µm at the capillaries, whereas the aggregate cross-sectional area increases from ∼1 cm^2^ to ∼10^3^ cm^2^ (1, 8). However, expanding the cross-sectional area of the daughter vessels can be costly because of the construction and maintenance of redundant channels. Murray’s law explains how the costs of running the vascular system can be minimized by controlling the diameter reduction ratio (*9, 10*). The same framework has been utilized for rationalizing the observed diameter reduction ratio in the xylem in plants, for which the constructing cost of conduits is the primary factor limiting the expansion of the cross-sectional area of the xylem vessels (*11*).

The airway system in a human lung exhibits a similar branching architecture. A single trachea bifurcates into ∽2^23^ terminal branches, where most alveoli that mediate gas exchange with blood capillaries are developed. In this bronchial network, the expansion of the aggregate cross-sectional area of the daughter channels is limited by the airway volume, in such a way that the air transport to the alveoli is maximized for a given amount of inhalation air (*12, 13*). Nevertheless, this rationale explains the airway branching only in conducting airways where no gas exchange occurs, as shown in Fig. 1.

**Fig. 1.**
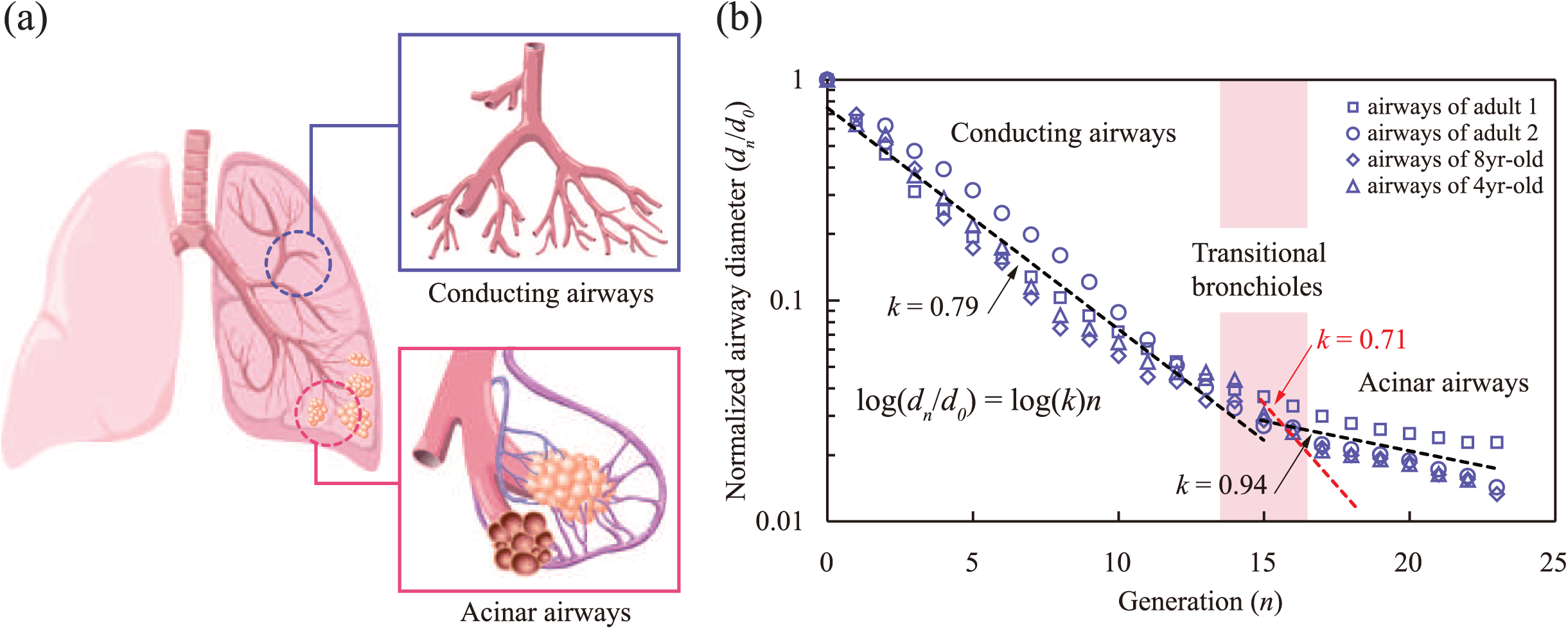
Anatomic observation of the airways of human lungs. (**A**) Schematic illustration of conducting airways (blue box) and acinar airways (red box). The hierarchical airway network consists of dichotomous trees with 23 generations. The transferred air diffuses to capillaries enclosed in the alveoli, most of which are attached to the 22nd and 23rd generations of the airways. (**B**) Reduction in normalized airway diameter along with airway generation. The diameter reduction ratio is 0.79 in the conducting airways, whereas it shifts to 0.94 in the acinar airways. Note that Murray’s law for diffusion in insects, *k* = 0.71, cannot explain the acinar airways reduction ratio (red dashed line). Data were taken from Finlay (*8*) and Weibel (*14*).

It has been supposed that the change in gas transfer mode from advection to diffusion is responsible for the transition in the reduction ratio of bronchial airways (10, 14, 15). Explicit analytical or computational studies, often combined with fractal modeling, focused on the velocity field in airways (*16*), permeability in acinus (*17*), breathing irregularity (*18*), asymmetric branching (*19-20*), and acinus morphology (*21*). The principle of the cost minimization for diffusive mass transfer was developed, which successfully provided the rationale for the spiracle pore networks of insects (*22, 23*). However, this model cannot be directly applied to acinar airways because the observed diameter reduction ratio of acinar airways (*k*=0.94) is much larger than that of Murray’s law for diffusion (*k*=0.71).

Here we present a model for the hitherto unexplained diameter reduction ratio in the acinar airways, *k*=0.94. The theory is experimentally supported by a microfluidic chip mimicking lung airways. With a simplified airway geometry that is amenable to mathematical analysis, our model captures an essential physical picture responsible for the observed diameter reduction ratio.

### Results

#### Theoretical Analysis

We begin with an analysis of the oxygen transfer in human lung airways. During a 4 s period of inhalation, a negative pressure in the pleural cavity induces the expansion of alveoli, and the total volume increase reaches approximately 500 ml (*24*). As a result of the increase in the cross-sectional area of the airways, the average speed of the air flow decreases from about 0.5 m/s in the trachea to 1 μm/s in the alveoli. The dominant oxygen transfer mechanism can be examined using the Peclet number, Pe = *ul/D*, the ratio of the advective to the diffusive mass transfer rates, where *l* is the length of a single airway branch (*l* ∼ 1 mm), *u* is the speed of the air flow, and *D* is the oxygen diffusion coefficient in air (*D* ∼ 0.2 cm^2^/s). Using the data of the cross-sectional area of airways for each generation (*8, 14*), one can find that Pe < 1 after transitional bronchioles, which suggests a shift in the dominant oxygen transfer mechanism from advection to diffusion (*10, 15*).

We develop a mathematical model of the oxygen transport in the acinar airways. Since the oxygen transfer through the channels via diffusion depends on their cross-sectional area, a trumpet model can be used (*25*). We construct a geometric model of the acinar airways as a stepwise channel with a unit depth by arranging all the channels side by side in such a way as to retain an equivalent aggregate cross-sectional area for each generation, as shown in Fig. 2. The stepwise channel is further simplified as a diverging duct enclosed by two curved boundaries. Assuming that the transition of diameter reduction ratio occurs near the 16th branch, we can express the cross-sectional area as *A*(*x*) = *A*_16_*e*^*x/a*^ with *a* = *l*/ln(2*k*^2^), where *x* is the distance from the 16th branch, *A*_16_ is the total cross-sectional area of the acinar airways at the 16th generation, and *k* is the diameter reduction ratio. We assume that oxygen transport from the air to capillaries occurs mainly at the airways of the 23rd generation where more than half of the alveolae lie (*17, 26-28*).

**Fig. 2.**
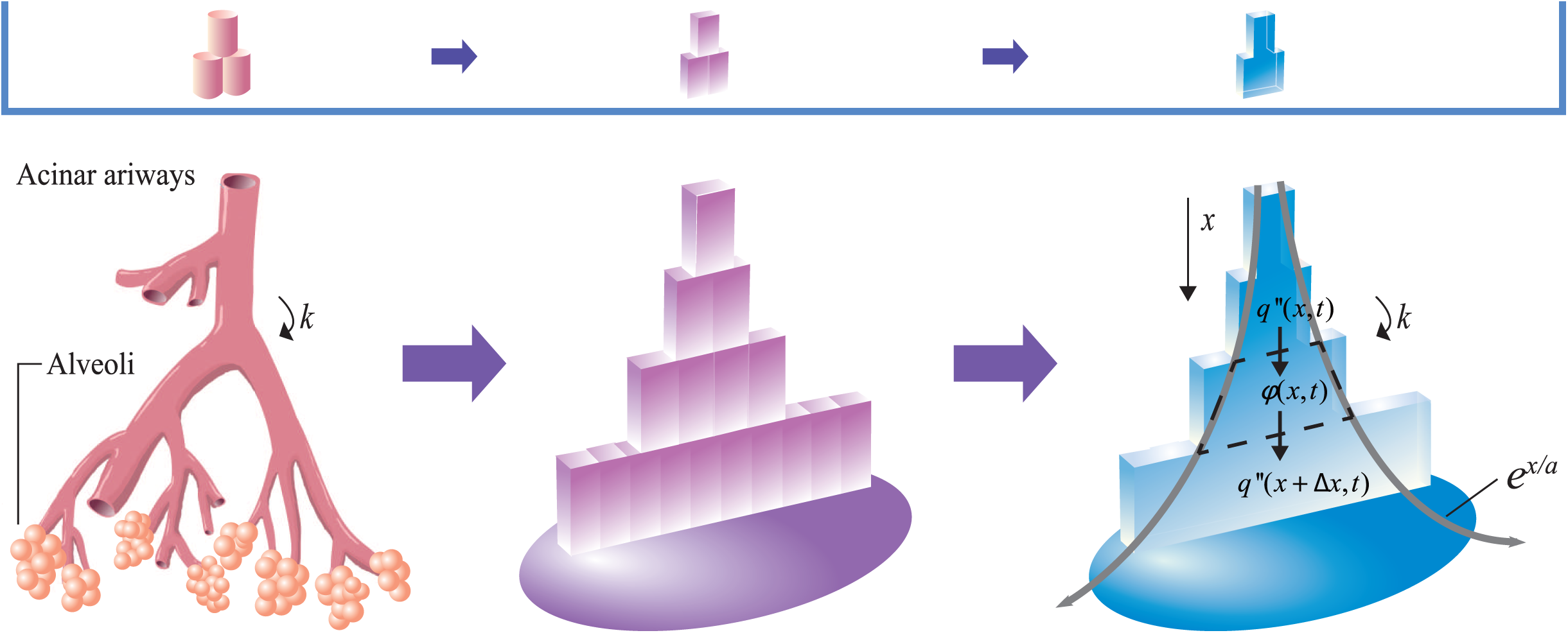
Schematic illustration of a trumpet model for acinar airways. The acinar airways consist of eight generations of airways and alveoli. The acinar airways can be assumed as a bundle of rectangular channels with the identical cross-sectional area on same generation. The sidewalls of the rectangular channels do not affect the vertical diffusion, so it can be assumed to be a single trumpet channel that expands like an exponential function involving the reduction ratio *k* and single channel length *l*.

We examine the oxygen conservation in an infinitesimal control volume, as shown in Fig. 2. Because sidewalls of a single airway prevent diffusion to neighboring airways, we only consider longitudinal diffusion along the *x*-axis, which can be formulated as

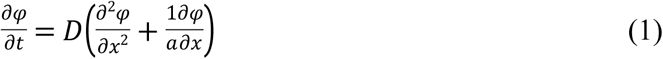

where *φ = φ*(*x, t*) is the partial pressure of oxygen in the acinar airways, and *t* is the time after inhalation begins. Solving Eq. (1) requires an initial condition and two boundary conditions. We assume that the oxygen partial pressure in the acinar airways is the same as that of the blood capillary when inhalation begins because an exhalation time of 4 s is greater than the diffusion time scale *L*^2^/(4*D*) ∼ 0.8 s, where *L* ∼ 8 mm is the diffusion length from the 16th to 23rd airways. Thus, the initial condition is written as *φ*(*x*, 0) = *φ*_*c*_, where *φ*_*c*_ is the oxygen partial pressure of the blood capillary. Once inhalation begins, fresh air is supplied from the conducting airways. Accordingly, we assume that the oxygen partial pressure at the inlet of the 16th airway branch remains the same as that in the fresh air during the inhalation. This leads us to write a boundary condition as *φ*(0,*t*) = *φ*_*a*_, where *φ*_*a*_ is the oxygen partial pressure in the fresh air. The other boundary condition comes from the continuity of the oxygen transfer in the alveoli, where oxygen diffuses to capillaries across the alveolar membrane,

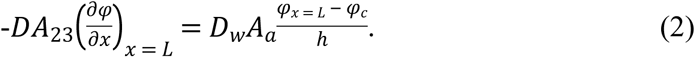

Here *A*_23_ is the total cross-sectional area of the acinar airways at the 23rd generation, *D*_*w*_ is the diffusion coefficient of oxygen in water, *A*_*a*_ is the total surface area of the alveoli, and *h* is the thickness of the alveolar membrane.

We obtain an analytical solution of equation (1) consisting of steady and transient parts (see Supplementary information). However, the oxygen partial pressure of the acinar airways becomes virtually steady within one-tenth of the inhalation duration, and so the transient part is negligible. The distribution of the oxygen partial pressure in the acinar airways is given by

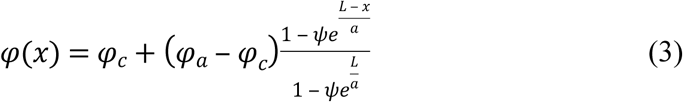

where *ψ* = (*aD*_*w*_*A*_*a*_)*/*(1−*hDA*_23_). Because *a* is a function of *k*, Eq. (3) allows us to infer the dependence of the oxygen transfer rate on the diameter reduction ratio. As shown in the inset of Fig. 3, the model predicts that the oxygen transfer rate increases with the diameter reduction ratio because of the expansion of the cross-sectional area.

**FIG. 3.**
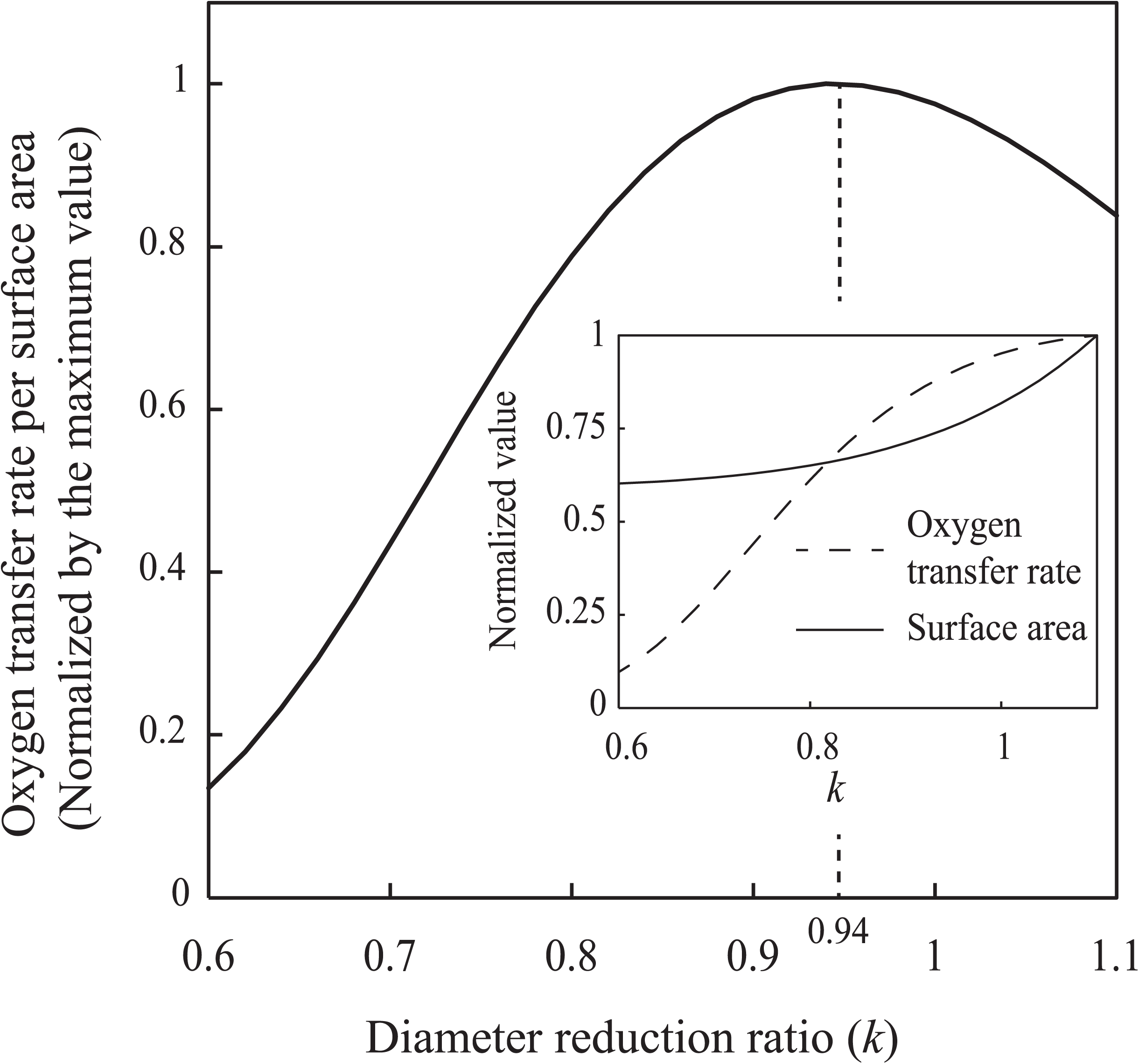
Oxygen transfer rate per surface area of acinar airways versus diameter reduction ratio. The oxygen transfer rate per surface area peaks at a diameter reduction ratio of 0.94, for which the energy cost for transporting a given amount of oxygen is minimized, provided that the energy investment is proportional to the surface area of acinar airways. The inset shows the dependence of oxygen transfer rate and surface area of acinar airways on the diameter reduction ratio. The values are normalized by their maximum values, respectively.

Acinar airways comprise a thin layer of epithelial cells covering airways and alveolae. Therefore, building an additional airway demands costs for construction and maintenance of the epithelial cell layer, which are proportional to the total surface area of the acinar airways. This reminds us of the fact that the expansion of the cross-sectional area in plant xylem vessels is limited by the carbon investment for the construction of the vessels (6). We thus suggest that the total surface area of the acinar airways is the primary limiting factor. Fig. 3 presents the oxygen transfer rate 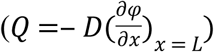 divided by the surface area of the acinar airways versus the diameter reduction ratio, revealing that *Q*/*A* is maximized when *k* is 0.94. We thus infer that a ratio of 0.94 is the optimal reduction ratio to minimize the cost of the acinar airways for oxygen transfer. This theoretical value is indeed consistent with the anatomic observations shown in Fig. 1b. On the basis of the model, it can be deduced that the transition of the diameter reduction ratios between the conducting (*k* = 0.79) and acinar (*k* = 0.94) airways saves more than 30% of the epithelial cell layers compared with the case without this transition. Consequently, our model suggests that this transition occurs for reducing energy consumption in the acinar airways.

To test the validity of our theory for the dependence of oxygen transfer rate on *k*, we carried out experiments using a microfluidic device mimicking an airway network in the lung (see Supplemental information). As shown in Fig. 4a, the microchip consists of two channels containing oxygen-rich air and oxygen-depleted water. Oxygen diffuses across a thin PDMS membrane from oxygen-rich air to oxygen depleted water stream, which was pumped into the channels by syringe pumps. Measuring the oxygen concentration at the outlet of the oxygen-depleted water channel for various oxygen-rich air channel dimensions yielded the oxygen transfer rates per surface area of the oxygen-rich channels as a function of *k*. As shown in Fig. 4b, our measurements are in a good agreement with our model prediction, providing an experimental support of the model.

**FIG. 4.**
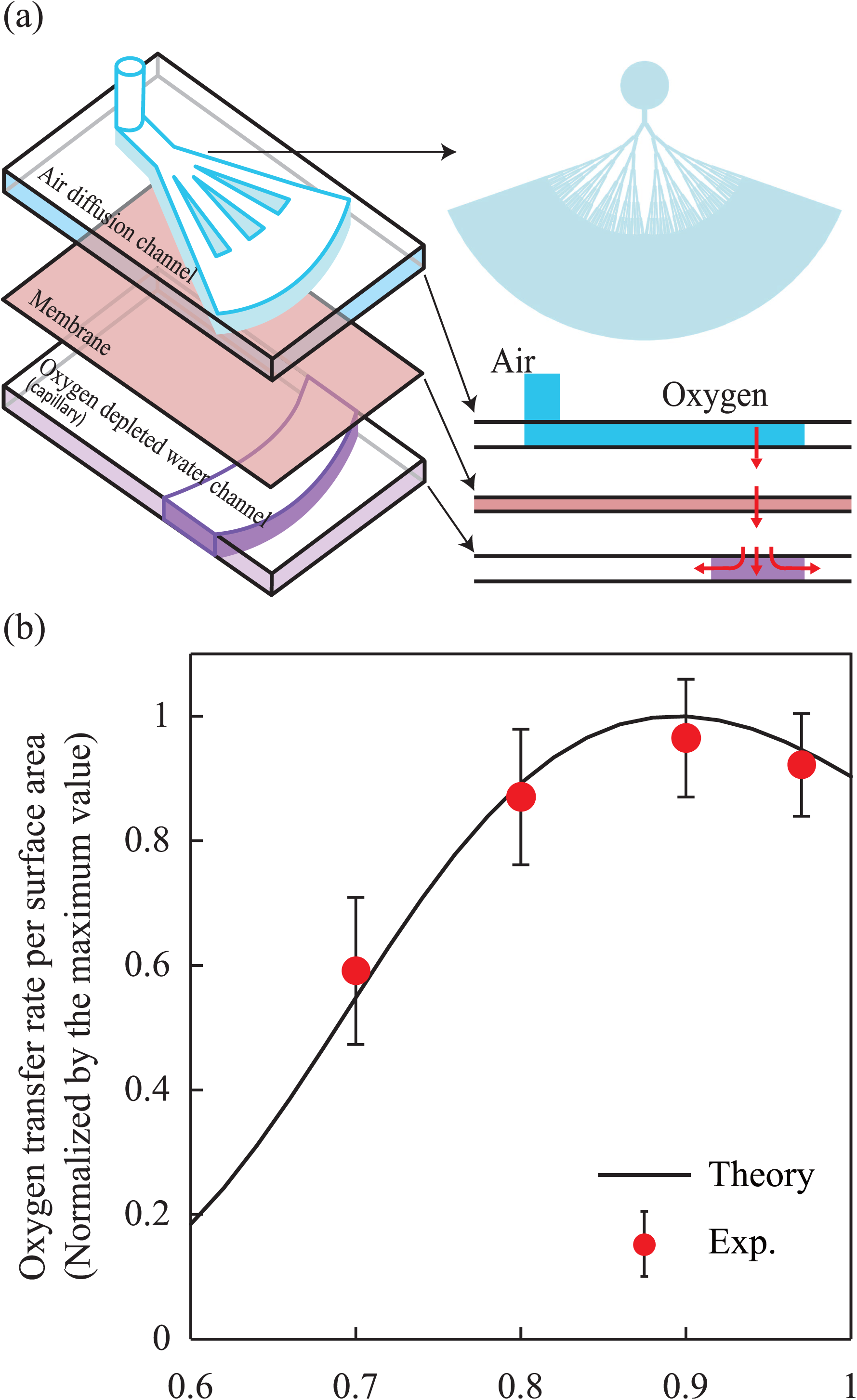
Measurements of oxygen transfer rates per surface area using a microfluidic chip mimicking acinar airways. (**A**) Schematic illustration of the lung-on-a-chip device constructed of PDMS. The microchip consists of two chambers containing oxygen-rich air and oxygen-depleted water, respectively, along with membrane between the two chambers. Each air diffusion channel branches with a specific diameter reduction ratio, *k*. The red arrows indicate the direction of the oxygen diffusion. (**B**) Dependence of the normalized oxygen diffusion rate per surface area on the diameter reduction ratio, *k*. The line and circles respectively correspond to the theoretical predictions and experimental results with different values of *k*. Error bars indicate the standard deviation resulting mainly from the sensing noise of the oxygen sensor.

### Discussion

Originally developed for blood vessels, Murray’s law describes branching structure for minimizing energy cost for convective transport and metabolism of blood (*9, 10*). This design principle of branching structure has been used as a framework to understand various natural network systems under the assumption that natural branching networks have evolved in such a way as to minimize the cost for transferring a given quantity of matter (6, 7). Murray’s law for conducting airways of lung explains the minimization of energy consumption for oxygen transport for a given volume of air inhalation (*8, 10, 17*). Murray’s law for plant xylem elucidates the minimization of the primary energetic cost caused by sap flow and vessel construction (*11*). Interestingly, despite various forms of energetic costs, Murray’s law for these networks commonly consider convection transport energy and vessel volume. The relation is mathematically expressed by an equation showing that the sum of cubes of the diameters of vessels in each generation is the same, yielding the diameter reduction ratio of 0.79. Murray’s law for diffusion can be formulated mathematically by replacing the energy cost for convective transport with the energy cost for diffusive transport. Then the sums of diameter squares in each generation are constant, leading to a diameter reduction ratio of 0.71 (22, 23).

However, the biological data of acinar airways in human lungs cannot be explained by either of the aforementioned models. In the present study, we elucidate the data by considering the amount of diffusive transport of oxygen per surface area of airways. This implies that the dominant cost for the construction and maintenance of acinar airways comes from epithelial cell layers of airways and alveolae, which should be measured by the surface area rather than the volume.

Although advection is typically cost effective on the organismal scale, diffusion is a rapid, reliable, and cheap way to transfer matter on the cell scale (*1, 29*). In mammals relying on air breathing, the characteristic diameter of the terminal branches of the airways is on the order of 100 μm, regardless of the body size (*14, 30, 31*). Thus, gas transport via diffusion would be more effective near the terminal branches. Accordingly, the transition of the oxygen transfer mode is also expected in the lung airways of other species, which may cause the transition of the diameter reduction ratios. Indeed, some anatomical data on the lung airways of other species, including rats, rabbits, and canines, show a transition in the diameter reduction ratio as in human lung airways (*30, 31*). This optimal strategy of acinar airways can guide design and construction of artificial networks where one needs to maximize fluid transport to a given area (*32-34*).

### Materials and Methods

#### Microfluidic chip mimicking acinar airways

The PDMS structure was prepared using a 10:1 mixture of Sylgard-184 (Dow Corning) cured by baking in a vacuum oven at 80°C for 30 min. The lung-on-a-chip device consisted of two layers separated by a PDMS membrane with a thickness of 40 μm. An eight-generation microchannel network for mimicking acinar airways was fabricated in the upper layer, and a one-way microchannel corresponding to blood capillaries was fabricated in the lower. The thicknesses of both layers were 100 μm, and the width of the first-generation channel was 300 μm. The inner wall of the microchannels was coated with 0.6 wt% Teflon-AF (DuPont 601S2 and 3M Fluorinert Electronic Liquid FC40) to prevent the oxygen from diffusing out of the microchannels. In the upper layer, oxygen diffused from one open end of the microchannel to the other closed end, where oxygen was transferred to the oxygen depletion channel in the lower layer through the membrane. In the oxygen depletion channel, the flow rate was controlled using a syringe pump (LSP 04- 1A, Baoding Longer Precision Pump), and sodium sulfite was used to remove the oxygen dissolved in the water. We produced four devices with the same membrane surface area (150 mm^2^) but different cross-sectional area reduction ratios of 0.7^2^, 0.8^2^, 0.9^2^, and 0.97^2^. We measured the oxygen concentration at the outlet of the lower layer microchannel using an oxygen microsensor (OX-100, Unisense), and then calculated the oxygen transfer rate.

## H2: Supplementary Materials

Section S1. Solution of Equation (1)

Table S1. Typical geometry of acinar airways (*7*).

Fig. S1. The distribution of oxygen partial pressure φ * obtained from transient and steady solutions.

Fig. S2. Transient and steady solutions of oxygen transfer rate per surface area versus diameter reduction ratio. Unsteady effects diminish in 1.28 s.

## Author contributions

K.P. carried out the experiments. K.P. and T.S. developed the mathematical model. K.P., Y.-J.C., N.L.J, W.K. and H.-Y.K. wrote the paper. W.K. and H.-Y.K. conceived and supervised the project.

## Data and materials availability

All data needed to evaluate the conclusions in the paper are present in the paper and/or the Supplementary Materials. Additional data related to this paper may be requested from the authors.

